# Combinatorial constraints predict that mitochondrial networks contain a large component

**DOI:** 10.64898/2026.03.25.714309

**Authors:** Raphael Mostov, Greyson R. Lewis, Moumita Das, Wallace F. Marshall

## Abstract

Mitochondria often form branching membrane networks distributed throughout the cell interior. In many, but not all, cell types, these networks are observed to consist of one large connected component together with many smaller fragments. Why does this pattern arise? Does it reflect a specific biological function, an external biophysical constraint, or something simpler? Using results from extremal graph theory, we prove a new theorem which suggests that, under a sufficiently broad sampling of the space of mitochondria-like graphs, the predominance of three-way junctions makes the appearance of a large component likely. This suggests that, in some settings, a large component may serve as a useful null model for mitochondrial network structure rather than requiring a dedicated explanation. More broadly, our result points towards testable predictions, since systematic deviations from this baseline may help reveal additional constraints or mechanisms shaping mitochondrial morphology.

## 1 Introduction

While often depicted as bean-shaped organelles, mitochondria frequently form branching networks distributed throughout the cell interior. Mitochondrial network morphology has been observed to vary with disease state [2], cell differentiation [3], cell-cycle transitions [4], and growth conditions [5]. Moreover, these network architectures are dynamic, constantly changing their connectivity [6], primarily through fission, in which a network branch splits, and fusion, in which two branches join together [7]. Additional morphological transformations, including the outgrowth of a branch from another branch and its inverse, resorption, have also been reported (summarized in [8]).

Despite conspicuous morphological differences across cell states, as well as dynamic rearrangement over time within a single cell, one feature appears repeatedly in mitochondrial morphology: networks often consist of one large connected component together with many smaller fragments. In the largest currently available data set on mitochondrial network morphology, comprising 2883 networks in *S. cerevisiae* cells analyzed by Viana et al. [1], 97% of networks have over half of their total length concentrated in one large component (see Figure 2C). Though less dramatic, similar trends have also been quantified in mouse embryonic fibroblasts [9, 10], adult guinea pig myocytes [11], and reported in at least six other cell types [12].

Not all cell types exhibit a single large component (see below). Nevertheless, the fact that a dominant large component appears so frequently across cell types calls for explanation.

One possible explanation is functional. Organizing the mitochondrial network into a single connected component may have important consequences for mitochondrial activity. For example, transport of metabolic intermediates could occur by diffusion through mitochondrial tubules, with transport efficiency depending on the level of connectivity [13]. Distribution and recombination of mitochondrial nucleoids would likewise be facilitated if mitochondria formed one connected compartment. Mitochondrial energy production depends on a proton gradient across the mitochondrial membrane, and fusion into a single large component could electrically couple a larger portion of the network. In some cases, increased mitochondrial connectivity has been linked to increased ATP production [14]. These arguments help explain why a large component might be advantageous, but they do not explain why such a structure should arise so often.

Given the impact of connectivity on function, one possibility is that cells have evolved a homeostatic mechanism that monitors mitochondrial activity and adjusts fission and fusion rates through feedback until some desired level of activity is reached. Alternatively, cells might directly monitor mitochondrial network structure and tune fission and fusion until a single connected component is achieved. However, such mechanisms remain speculative, raising the question of whether a simpler explanation, not involving explicit homeostatic regulation, might be sufficient.

Another possibility is that the balance of fission and fusion lies near a critical point, so that slight perturbations in these rates can cause a network to transition rapidly into a large component [15, 16]. Consistent with this idea, Zamponi et al. found that component-size ratios in mouse embryonic fibroblasts were compatible with such models [10]. On this view, the frequent appearance of a large component could reflect the tuning of mitochondrial dynamical processes by natural selection. More generally, many factors could shift the effective balance of fission and fusion, including enzyme kinetics, spatial confinement, and organelle packing, all of which may alter the probability that nearby branches encounter one another and fuse.

Such models remove the need for explicit feedback control over connectivity, but they still rely on specific tuning of mitochondrial dynamics to produce a large component. But what if there is an even simpler explanation—one that is independent of the detailed rates of mitochondrial dynamics, and perhaps even independent of any specific biological function?

In this paper, we apply results from extremal graph theory to hypothesize such an explanation. We prove a new theorem suggesting that if a distribution of mitochondrial networks samples broadly from the space of admissible mitochondria-like graphs, then the predominance of three-way junctions makes the presence of a large component likely. In this sense, the frequently observed large component may, in some settings, be understood as a useful null model for mitochondrial network structure rather than as a feature that necessarily requires explanation. Our aim is not to deny the importance of biological mechanism, but to identify a baseline against which additional constraints and functions can be more clearly interpreted.

## 2 Results

### 2.1 Graph representation of mitochondrial networks

Our first task is to define a useful graph-theoretic representation of mitochondrial network morphology. A natural approach is to represent a mitochondrial network as a graph whose vertices are connected by edges (for an introduction to graph theory in the context of mitochondrial networks, see [8]). To convert an image of a mitochondrial network (for example, Figure 1A) into a graph (Figure 1B), we assign a vertex to each branch tip and to each three-way junction. This leads to the following definition, which will be used throughout the remainder of the paper:

**Figure 1.**
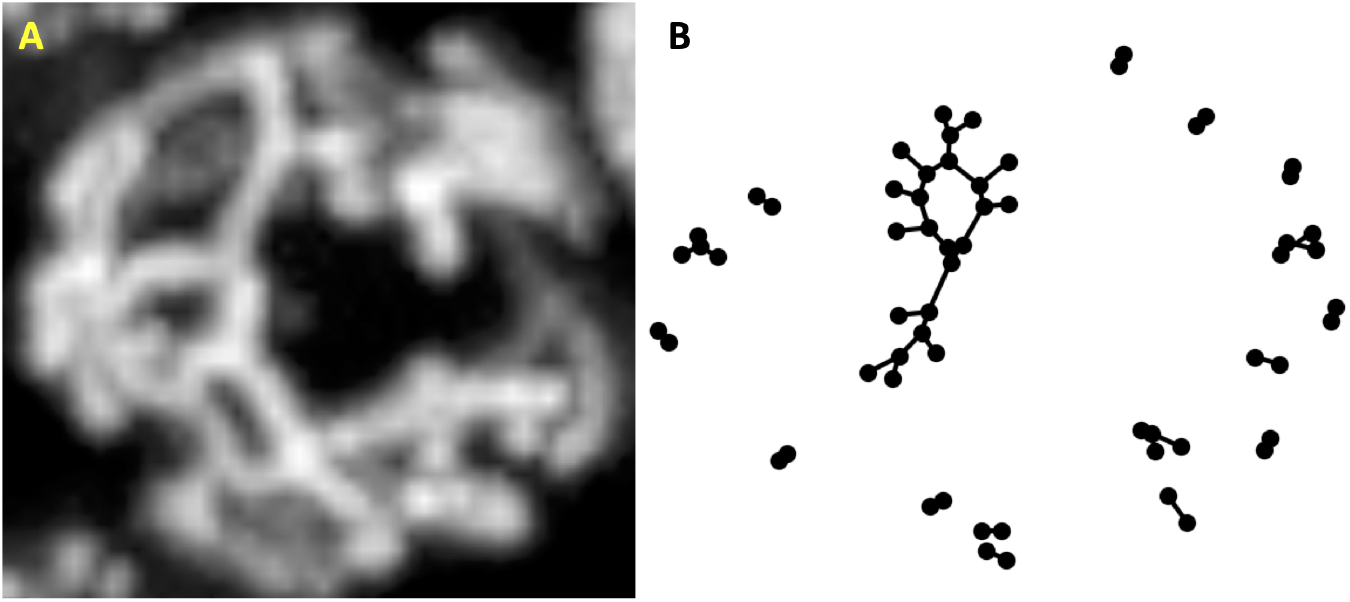
Graph representation of a mitochondrial network. (A) Max-intensity projection of a 3D image of a budding yeast cell expressing mito-dsRed, grown in glycerol and imaged by spinning-disk confocal microscopy. (B) Mitochondrial graph structure of the yeast cell in glycerol computed using MitoGraph 2 software [1].

#### Definition 1

. A *mitochondria graph G* is a simple unlabeled graph containing exclusively degree-one and degree-three vertices. We denote by *N* the total number of vertices, of which *n* have degree three and *N* − *n* have degree one.

This is not the only possible graph representation of mitochondrial networks. Because mitochondrial networks are physical structures embedded in Euclidean space, with both surface area and volume, most previous work has either assigned lengths to edges or subdivided edges into chains of degree-two vertices so that each original edge is represented by a sequence of unit-length segments [1, 16]. However, to establish a more general result, we do not use these more detailed representations here. Instead, we justify our choice empirically: the Pearson correlation coefficient between the number of vertices in a connected component and its total length is 0.9757 in the Viana et al. dataset [1] and 0.9460 in the Holt et al. dataset [15].

Finally, real mitochondrial networks are not always perfectly constrained to have only degree-one and degree-three vertices, nor to have at most one edge between any pair of vertices. Multiple edges between the same pair of vertices appear in the Viana et al. dataset only 3.7% of the time, which is rare enough for us to ignore for present purposes. Degree-four vertices are also observed, but only about 7.3% of the time, and time-series data from live cells suggest that these vertices are transient, typically resolving into two degree-three vertices, likely due to mechanical instability [8].

### 2.2 Distribution of component sizes supports a single large component in budding yeast mitochondrial networks

Using the graph representation above, we first analyze existing data to determine whether mitochondrial networks are in fact dominated by a single large component. In the Viana et al. dataset, shown in Figure 2A and Figure 2C, budding yeast cells exhibit a clear pattern: most vertices and most total network length are concentrated in a single large component. Moreover, Figure 2G shows that the second-largest connected component in budding yeast cells contains markedly fewer vertices than the largest component.

**Figure 2.**
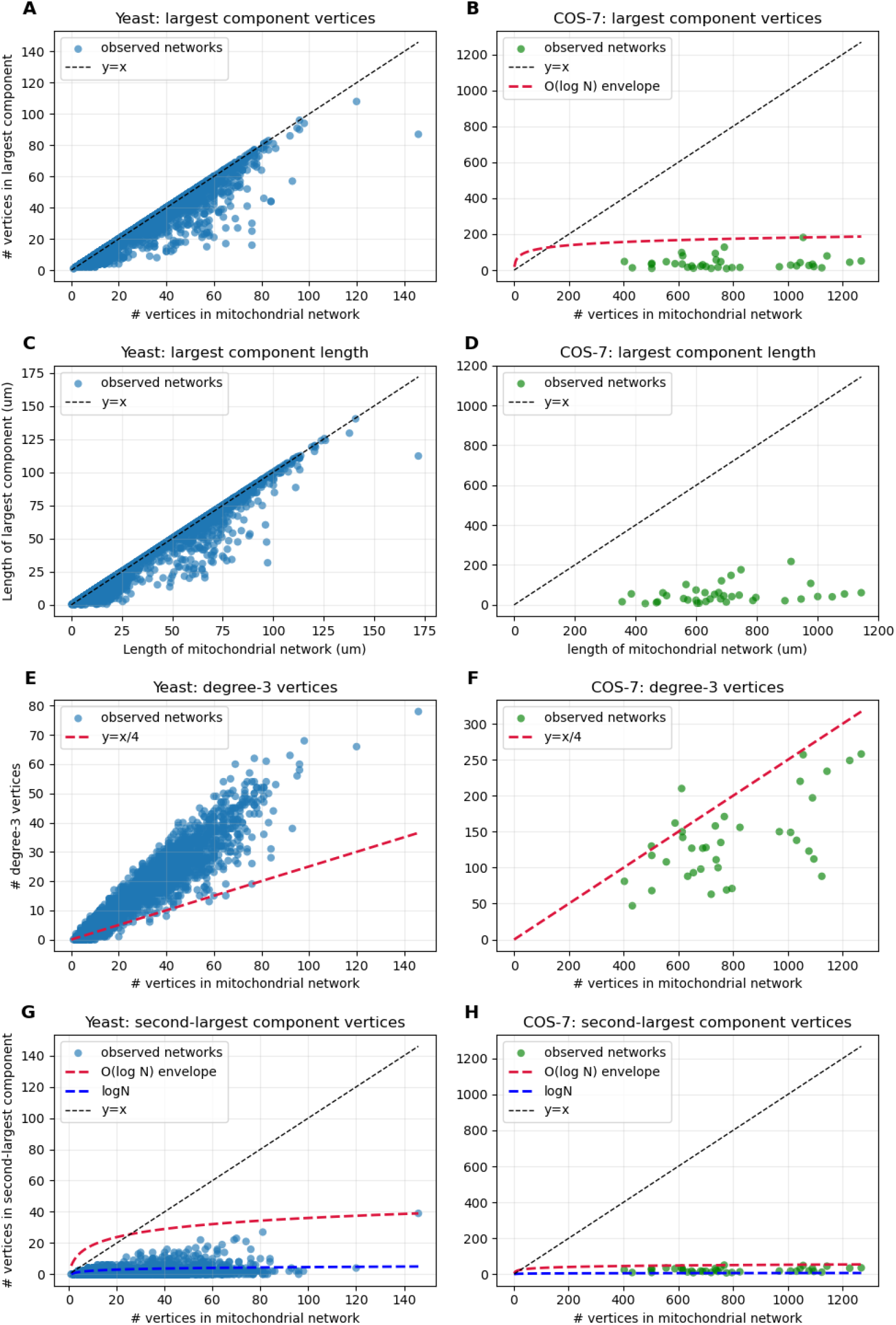
(A) Number of vertices in the largest network component plotted against the total number of vertices, *N*, in yeast cells; (B) in COS7 cells. (C) Length of the largest component plotted against total length in yeast; (D) in COS7. (E) Number of degree-three vertices, *n*, plotted against *N*, together with the line *n* = *N/*4, in yeast; (F) in COS7. (G) Number of vertices in the second-largest network component plotted against *N*, together with log *N*, in yeast; (H) in COS7. Budding yeast data are from [1] (*S* = 2883), and COS7 data are from [15] (*S* = 37).

A similar analysis of mitochondrial networks in COS7 cells, using data from Holt et al. [15], indicates that this pattern does not hold in that cell type. There, the mitochondrial network is not dominated by a single large component. We return to such cases in the Discussion. For now, we focus on explaining why mitochondrial graphs in many cell types are nevertheless dominated by a single large component.

### 2.3 Probability of a mitochondria graph having a single large component

We now have the context needed to state our main result.

#### Definition 2

Let *G* be a graph sampled from a probability distribution over graphs with *N* vertices. Then *G*_*L*_ denotes the event that *G* has a connected component with *O*(*N*) vertices and every other connected component has at most *O*(log *N*) vertices.

#### Theorem 3

*Let G be a mitochondria graph sampled uniformly from the set of mitochondria graphs with N vertices. Then* Pr(𝒢_*L*_) → 1 *as N* → ∞.

This theorem states that, for sufficiently large *N*, a uniformly sampled mitochondria graph will with high probability consist of a single large component containing *O*(*N*) vertices, while all remaining components have size at most *O*(log *N*). It is therefore a statement about the typical structure of large mitochondria graphs under uniform sampling, not a deterministic guarantee about any particular graph.

The theorem makes two predictions: first, that the largest connected component should contain *O*(*N*) vertices, and second, that the second-largest connected component should contain at most *O*(log *N*) vertices. As shown in Figure 2A and Figure 2G, these predictions are broadly consistent with the Viana et al. dataset [1].

The proof is given in the Methods. A key assumption, stated explicitly in the theorem and used throughout the proof, is that graphs are sampled uniformly; that is, every mitochondria graph with *N* vertices is equally likely to be chosen.

A natural question is whether real mitochondrial networks are in fact sampled uniformly from this space. We do not claim that they are. Rather, the value of the uniform model is that it identifies the prevalence of large-component graphs within the full combinatorial space of mitochondria graphs with *N* vertices. In this sense, the theorem shows that large components are highly prevalent in the underlying graph space and may therefore be expected under empirical distributions that sample that space sufficiently broadly. We therefore propose broad sampling of the admissible mitochondria-graph space as a useful null model for the frequent appearance of a large component.

### 2.4 The null model is falsifiable

What about cell types in which this pattern is not observed? While we leave the broader explanation for such cases to the Discussion, the same framework also yields a prediction for heavily fragmented mitochondrial networks. Specifically, we have the following result:

#### Proposition 4

*Let G be a mitochondria graph sampled uniformly from the set of all mitochondria graphs with N vertices satisfying n* ≤ *N/*4, *where n is the number of degree-three vertices. Then the probability that no connected component of G has more than O*(log *N*) *vertices approaches* 1 *as N approaches infinity*.

This proposition implies that heavily fragmented mitochondrial networks should be expected to satisfy *n* ≤ *N/*4. Consistent with this prediction, 86.5% of the COS7 networks satisfy *n* ≤ *N/*4.

This also suggests a clear way in which the null model could fail. If one were to observe a distribution of mitochondrial graphs in which *n* ≤ *N/*4 and yet a large component is still typically present, or conversely a distribution in which *n > N/*4 and no large component is typically present, then the explanation developed here would be insufficient for that case, and an additional explanation would be required. Indeed, many such reasons are possible; for example, a biological distribution may fail to sample broadly from the space of mitochondria graphs because of geometric, dynamical, or functional constraints (see Discussion).

## 3 Method (Proof of Theorem)

### 3.1 Outline of Proof

We remind the reader that from Definition 1, a mitochondria graph is a simple unlabeled graph with *n* degree-three vertices and *N* − *n* degree-one vertices, and, from Definition 2, that *G*_*L*_ is the event in which a graph has a connected component with *O*(*N*) vertices, and no other components with more than *O*(log *N*) vertices.

Now, suppose *G* is a graph sampled uniformly from the set of mitochondria graphs with *N* vertices. To prove Theorem 3, we must show that

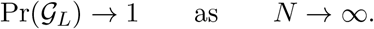

To do this, we will first show:

#### Proposition 5

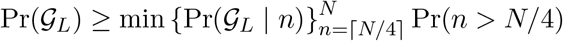

If we then show that both terms in the lower bound of Pr(𝒢_*L*_) approach 1 as *N* approaches infinity, then this will immediately imply Theorem 3. We will apply an extremal graph theory result from Molloy and Reed [17] to show that:

#### Proposition 6

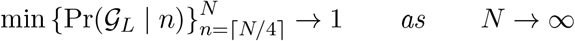

In doing so, we will see that Proposition 4 emerges as a corollary. Next, we will show:

#### Proposition 7

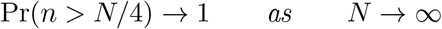

This will be done by first applying the Bender-Canfield formula [18], which asymptotically approximates the number of labeled graphs with a given degree sequence, to find upper and lower bounds on the number of mitochondria graphs *U*_*N*_ (*n*) with *N* vertices and *n* degree-three vertices:

#### Lemma 8

*Let U*_*N*_ (*n*) *be the number of mitochondria graphs with N vertices and n degree-three vertices. Then for large N*,

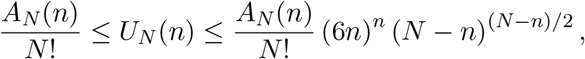

*where*

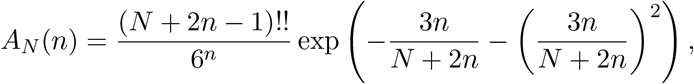

*and the double factorial denotes*

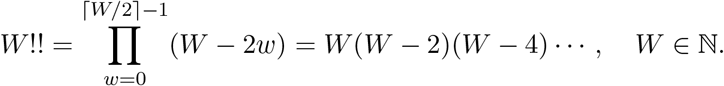

Using these bounds, we will then find a lower bound on the fraction of mitochondria graphs with *N* vertices satisfying *n > N/*4,

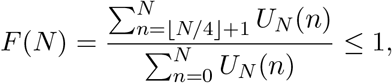

and then show that the lower bound approaches 1 as *N* goes to infinity, thereby showing that *F* (*N*) → 1 as *N* → ∞, and, subsequently, since *F* (*N*) = Pr(*n > N/*4) under the uniform distribution, showing Proposition 7. This will then complete our proof of Theorem 3.

### 3.2 Proof of Propositions 4, 5 and 6

We begin by showing Proposition 5:

*Proof of Proposition 5*. Let *G* be a mitochondria graph sampled uniformly from the set of mitochondria graphs with *N* vertices. Then,

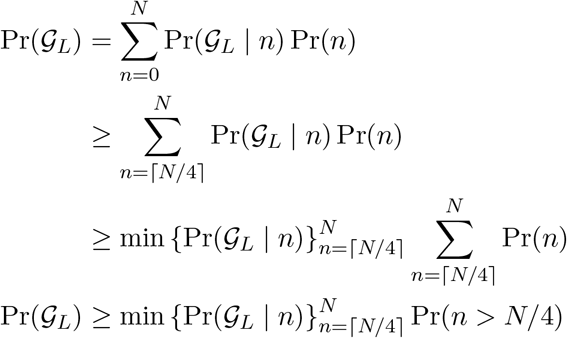

Next, we show Proposition 6. To do this, we will first need the following result form Molloy and Reed [17]:

#### Proposition 9

*Let λ*_0_, *λ*_1_, … *be nonnegative real numbers summing to* 1, *and consider random graphs having λ*_*i*_*N vertices of degree i, where each graph with that degree sequence is equally likely*.

- *If* ∑_*i*_ *i*(*i* − 2)*λ*_*i*_ *>* 0, *then* Pr(𝒢_*L*_) → 1 *as N* → ∞.
- *If* ∑_*i*_ *i*(*i* − 2)*λ*_*i*_ *<* 0, *then* Pr(𝒢_*S*_) → 1 *as N* → ∞, *where G*_*S*_ *is the event that a graph has no connected components with more than O*(log *N*) *vertices*.

We will then apply the Molloy Reed condition to a sequence of mitochondria graphs in order to show Proposition 6:

*Proof of Proposition 6*. For mitochondria graphs, the Molloy Reed constant is

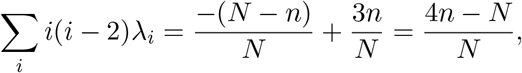

and so

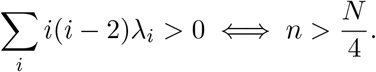

Let 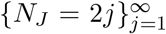 be the sequence of positive even integers, and let *G*_*j*_ be a mitochondria graph sampled uniformly from the set of mitochondria graphs with *N*_*j*_ vertices (by the handshaking Lemma, the total degree of a graph must be even, and mitochondria graphs contain only vertices of odd degree). Define the sequence 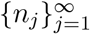 where *n*_*j*_*/N*_*j*_ = *λ*_3_ is approximately constant, and *n*_*j*_ *> N*_*j*_*/*4. If we condition on *n*_*j*_, then *G*_*j*_ becomes sampled from the distribution where each graph with *N*_*j*_ vertices, and *n*_*j*_ degree-three vertices, is equally likely. Thus, by Proposition 9, Pr(𝒢_ℒ_ | *n*_*j*_) → 1 as *N*_*j*_ → ∞. Subsequently, since we can choose *λ*_3_ to be any constant in which *n*_*j*_ *> N*_*j*_*/*4, then min 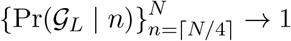 as *N* → ∞.

By instead defining *n*_*j*_ to satisfy *n*_*j*_ ≤ *N*_*j*_*/*4 in our proof of Proposition 6, we see that Proposition 4 emerges as a corollary.

### 3.3 Proof of Proposition 7

In order to show Proposition 7, we first need to derive upper and lower bounds on the number of mitochondria graphs *U*_*N*_ (*n*) with *N* vertices and *n* degree vertices from Lemma 8,

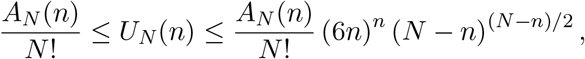

where

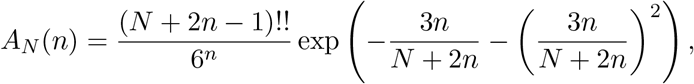

*Proof of Lemma 8*. The Bender-Canfield formula [18] asymptotically approximates the number of labeled graphs of a given degree sequence **d**. We use the version of this formula provided by Kawamoto [19]:

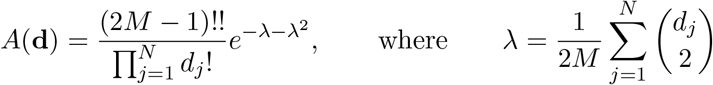

Here, *d*_*j*_ is the degree of vertex *j* and 2*M* = ∑*d*_*j*_. Applying this formula to a mitochondria graph with *n* degree-three vertices and *N* − *n* degree-one vertices, we get the desired *A*_*N*_ (*n*).

If *G* is a mitochondria graph with *N* vertices, then there are 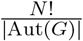 possible labelings of *G*. Here, |Aut(*G*)| is the number of possible automorphisms on *G*, where an automorphism *σ* is a permutation of the vertex labels of *G* in which *σ*(*u*), *σ*(*v*) form an edge if and only if vertices *u* and *v* form an edge. Let *k* ∈ ℕ^+^ index the set of mitochondria graphs with *N* vertices and *n* degree-three vertices, then for large *N*,

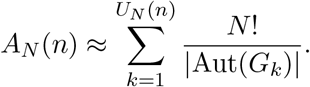

Since 1 ≤ |Aut(*G*_*k*_)| for all *k*, then for large *N*,

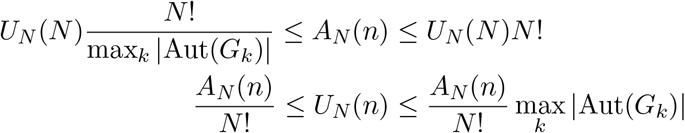

We thus seek to find an upper bound for max_*k*_ |Aut(*G*_*k*_)|. A degree-one vertex in *G* is either connected to a degree-three vertex, or another degree-one vertex thus forming a subgraph isomorphic to *K*_2_. Write *G* = *H* ⊔ *rK*_2_, where *r* is the number of *K*_2_ subgraphs in *G*. Since no vertex in *rK*_2_ is connected to a vertex in *H*, then Aut(*G*) ≌ Aut(*H*) *×* Aut(*rK*_2_), and so,

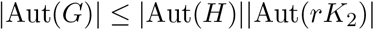

It is known that |Aut(*rK*_2_)| = 2^*r*^*r*!, and there are at most (*N* − *n*)*/*2 copies of *K*_2_ in *G*, and so

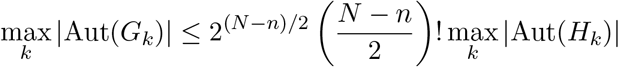

Turning our attention to the subgraph *H*, the labels of the *n* degree-three vertices can be permuted at most *n*! different ways, and each degree-three vertex is connected to at most three degree-one vertices whose labels can be permuted at most 3! = 6 different ways. Thus,

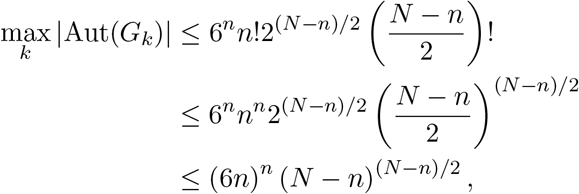

giving us the desired upper bound.

Now, we can prove Proposition 7:

*Proof of Proposition 7*. Let *G* be a mitochondria graph sampled uniformly from the set of mitochondria graphs with *N* vertices. Then

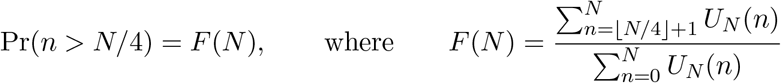

is the fraction of mitochondria graphs with *N* vertices satisfying *n > N/*4. It is therefore sufficient to show that *F* (*N*) → 1 as *N* → ∞. Write

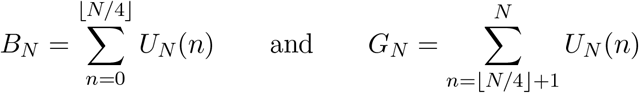

so that

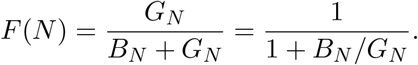

Since 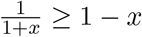 for *x* ≥ 0, then 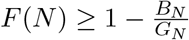. Since *F* (*N*) ≤ 1, then we need only show that 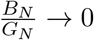 as *N* → ∞. Since 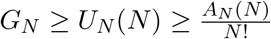, then by Lemma 8,

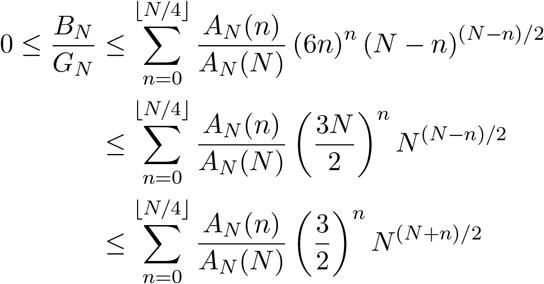

Next, we claim that

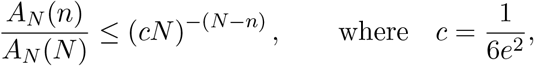

for all *n* ≤ *N*. To see this, write

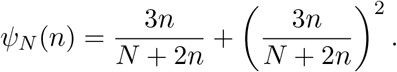

Then

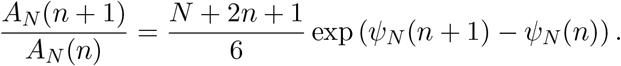

Since *n* ≤ *N*, then 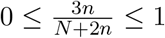, so 0 ≤ *ψ*_*N*_ (*n*) ≤ 2. Therefore, 0 ≤ *ψ*_*N*_ (*n* + 1) − *ψ*_*N*_ (*n*) ≤ 2, and so

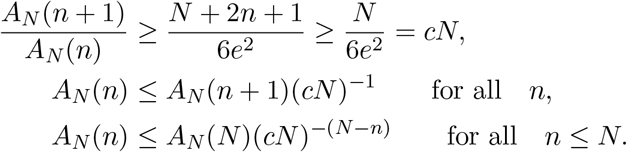

Therefore,

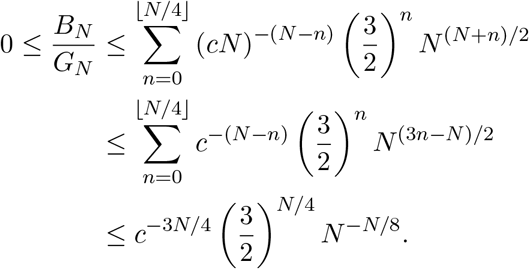

We see that 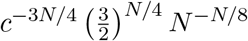 is dominated by the *N* ^−*N/*8^ term, which goes to 0 as desired.

## 4 Discussion

### 4.1 Dependence of our results on features of mitochondrial graphs

Our results indicate that a large component arises “for free” as a consequence of extremal graph theory, given the fact that mitochondrial graphs consist exclusively of vertices of degree 1 or 3. In some sense, though, this explanation just shifts the question to a different question, namely, why do mitochondrial graphs typically have this degree distribution? Degree 2 is impossible just by definition. Degree 4 or higher is presumably unstable due to the fluid dynamics of the mitochondrial membrane, but this is a question that deserves to be tested by direct experiments. In any case, we can say that while extremal graph theory explains the phenomenon for free, it only does so because biological or physical considerations have lead to mitochondrial graphs having the specific degree distribution that they have.

### 4.2 Why don’t all cells show mitochondria with a single large component?

We note that the presence of a single large component is not universal across all eukaryotes. Given the generality of the derivation above, why do some cells *not* show a single large component?

One possibility is that the abundance of disconnected components may simply be illusory. In many mammalian cells grown in culture, most of the mitochondrial material is clustered densely around the nucleus, where it has ready access to mRNA which is produced in the nucleus, while the rest is spread out more sparsely throughout the rest of the cell body, in order to deliver ATP to the cell’s extremity. It is much easier to image mitochondria in the flat regions of the cell away from the nucleus, where the mitochondria are sparser, due to the fact that two dimensional images can capture all of the material in a single focal. In contrast, the peri-nuclear region, which is where we would expect to actually see the large component if there is one, is much harder to visualize due to the limited axial resolution of light microscopy. Thus, it is crucial to reconstruct the full 3D mitochondrial network in order to decide if a single component is, or is not, present.

Setting aside cases where the lack of a large component is just due to incomplete imaging, there remain clear cases where the full network has been reconstructed and shown not to be dominated by a single large component. The largest data set for which we could find a counter example shows that across 37 different COS 7 cells, none of them have a component that makes up more than 8% of their total length [15](see Figure 2D). How can such cases be possible, in light of the highly generic nature of the results from extremal graph theory?

We have shown above that if the number of three-way junctions is small, then a single large component is unlikely, and this criterion is in fact met by the COS 7 cell data. In that sense extremal graph theory can also explain the cases in which a large component is not present. However, this explanation hinges on a specific feature of the graphs, namely a low number of three-way junctions, and that feature, itself, may require an explanation. why would some mitochondrial graphs have fewer three way junctions that others?

One likely explanation is that additional forces are being exerted on the mitochondrial components to push them into geometric configurations that are not consistent with the single component. Consider the example described above, in which a substantial fraction of the mitochondrial material is clustered around the nucleus and the rest distributed over the whole cell. Given a fixed quantity of mitochondrial material, it may simply be physically impossible to arrange a single component in this configuration. The single component would need to be relatively compact, and most likely to surround the nucleus and therefore unable to reach the rest of the cell body without fragmenting. This difference in organization between central and peripheral regions is likely generated by motor proteins pulling mitochondria on microtubules organized by a per-nuclear centrosome [20].

This specific example illustrates that mitochondria are not abstract graphs but physical networks, and subject to physical forces, for example those generated by the cytoskeleton, as well as to biological constraints such as a limited total quantity of mitochondrial membrane. It is the combination of mathematical structure, biological activities, and physical forces, that determine how the mitochondrial graphs are embedded in 3D space. It need not be the case that every possible graph is consistent with the constraints on physical embedding, and this could explain why classes of graphs that are predicted to be highly likely based on extremal graph theory might in fact not occur in physical mitochondria.

These hypothesized explanations, together with our result, points toward future testable predictions. As physical constraints are removed, for example by mutating linkers that couple mitochondria to the cytoskeleton, or changing cell shape to reduce the distance from the nucleus to the cortex, does a single large component start to emerge in cells that normally would lack it?

### 4.3 What is the expected size of of the large component?

So far, we have described conditions for a single large component where we define large relative to the rest of the components. But we have not directly stated what the expected size of the giant connected component that is likely to appear in a mitochondria graph is. Molloy and Reed do provide a formula that can, in theory, be used to find this quantity [21]:

#### Proposition 10

*(Molloy and Reed)*. *Take as given the premise of Proposition 9. The number of vertices in the large component is then*

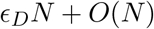

*Where*

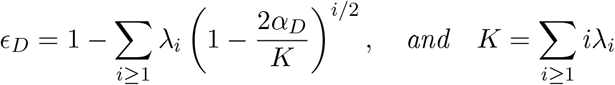

*and α*_*D*_ *is the smallest real positive α satisfying*

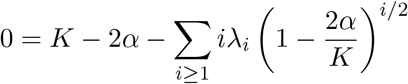

Given that the largest component cannot have more than *N* vertices and the expected size of the largest component has a linear error term, as well as the difficulty in generalizing this quantity to mitochondria graphs with *N* total vertices, we leave the search for the expected size of the largest component to future work. For similar reasons, we also leave finding a mathematical estimate of the probability that there is a single large connected component to future work.

### 4.4 Extremal graph theory as a tool for investigating cell biology

Mitochondria are not the only structures in the cell that can be described using graphs. The endoplasmic reticulum is the other obvious example, again consisting of a network of membrane tubes. Other examples include the actin cortex. Even chromatin inside the nucleus can be viewed as a network given the extensive set of contacts between different chromosomal regions revealed by modern methods like Hi-C. Any time there are large networks, there is the potential to apply extremal graph theory to predict statistically likely features of the network.

The presence of a single component is just one example. Another type of extremal property is the presence of sub-graphs with specific structures, such as K graphs or cycles. Analysis of such “network motifs” has been extensively employed in the study of cell signaling networks [22]. An extensive body of mathematics, including the famous theorems of Mantel and Turán, give conditions under which specific types of sub-graphs are guaranteed to arise in sufficiently large graphs. We believe that a productive line of investigation could be to search for network motifs in mitochondria and other physical cellular networks and then to ask whether those motifs can be explained purely by extremal graph theory. To our knowledge, network motif analysis has not been explicitly carried out for mitochondrial networks of the type considered here.

### 4.5 Conclusion – the large component as a null hypothesis

The results of this paper suggest that, under the assumptions considered here, a large connected component may be a reasonable null hypothesis for mitochondrial network structure. In our model, such a component becomes likely as the number of vertices grows, due to combinatorial features of the space of admissible mitochondria graphs. This should not be taken to mean that observed mitochondrial morphology is determined by combinatorics alone, nor that biological function, geometry, or network dynamics play no role. Rather, the point is that the frequent appearance of a large component may not by itself require a specialized explanation. Instead, it may be useful to view the behavior described here as a baseline, so that systematic deviations from it can help identify additional constraints or mechanisms relevant to mitochondrial organization.

## 5 Acknowledgments

We thank the late Clifford Marshall, Gav Sturm, Mary Mirvis, and Ximena Garcia Arceo for informative discussions about mitochondria and graph theory. This work was supported by NIH grant R35 GM130327 and by an External Research Project grant from the National Institute of Theory and Mathematics in Biology grant P0585526.

